# Mapping the magnetoreceptive brain: A 3D digital atlas of the migratory bird Eurasian blackcap (*Sylvia atricapilla*)

**DOI:** 10.1101/2025.03.04.641293

**Authors:** Nikoloz Sirmpilatze, Alessandro Felder, Dinora Abdulazhanova, Leonard Schwigon, Katrin Haase, Isabelle Musielak, Troy W. Margrie, Henrik Mouritsen, Dominik Heyers, Adam L. Tyson, Simon Weiler

**Author notes:** co-first authors (Nikoloz Sirmpilatze, Alessandro Felder, Dinora Abdulazhanova). co-supervisors (Dominik Heyers, Adam L. Tyson, Simon Weiler). corresponding authors*, **Corresponding author** D Heyers, AL Tyson, S Weiler.

## Abstract

Birds undisputedly range amongst nature’s foremost navigators. To successfully navigate between breeding and wintering quarters, they, in addition to other natural orientation cues, rely on their ability to sense the Earth’s magnetic field. For this reason, migratory birds have become key model species for studying the sensory mechanisms underlying magnetic field-guided navigation, as evidenced by the identification of several brain regions believed to be involved in processing magnetic field information. However, there is as yet no readily accessible, high-resolution three-dimensional (3D) brain atlas to serve as a common reference within and across studies. Here we provide the neuroscience research community with the first freely available, digital, high-resolution (25 *µm*), 3D bird brain atlas. It is based on light microscopy images from ten Eurasian blackcaps (*Sylvia atricapilla*), a night-migratory songbird widely used model species in magnetoreception and navigation research. We outline the individual steps for the creation of a brain atlas, from whole-brain imaging using serial-section, two-photon tomography, to the creation of an average template at an isotropic 25-*µm* voxel size, and finally to brain area segmentation and annotation. In this first version of the atlas, we have mapped a total of **24** brain areas, including **6** principal compartments, **13** conspicuous anatomical subdivisions common to all bird species and **5** functionally defined areas of the visual and trigeminal sensory systems involved in processing magnetic field information. This atlas is accessible via the standardised BrainGlobe Atlas API, making it compatible with a growing suite of computational neuroanatomy tools provided by the BrainGlobe Initiative. This integration enables precise alignment of future experimental data to a common coordinate space, facilitating collaboration, data visualization and sharing. Furthermore, this resource enables the accurate localization and comparison of implanted devices, injection sites, and/or cell populations across individual brains, both within and across studies.

## 1 Introduction

The evolutionary divergence of the avian brain, at least in part, reflects the unique morphological and physiological specializations required for successful procreation (Güntürkün and Bugnyar 2016; Stacho *et al*. 2020; Niu *et al*. 2022). Some bird species exhibit unique natural behaviors and cognitive specializations neither present in mammals nor in any other animal models and therefore offer an opportunity to gain insight into how the nervous system has adopted specialized sensory processing functions beyond what most mammals and humans might be capable of detecting. One advantage of flight over other forms of locomotion is the ability to cover vast distances. For example, migratory birds travel thousands of kilometers, often during the night and without the guidance of conspecifics, between wintering and breeding grounds. After they have completed their return migration once, many species achieve centimetres precision over distances of thousands of kilometres (Mouritsen 2018). In addition to using e.g. celestial orientation cues, these extraordinary navigational capabilities rely on the ability to sense the Earth’s magnetic field (Wiltschko and Wiltschko 1972; Mouritsen 2018; Wynn *et al*. 2022). However, despite major progress in understanding the sensory mechanisms underlying detection of this navigational cue (Zapka *et al*. 2009; Zapka *et al*. 2010; Hore and Mouritsen 2016; Xu *et al*. 2021; Görtemaker *et al*. 2022; Leberecht *et al*. 2023), magnetoreception remains one of the most enigmatic forms of sensation in systems neuroscience research (Mouritsen 2018; Nimpf and Keays 2022).

One of the most studied bird species in magnetoreception research in recent years is the common Eurasian blackcap (*Sylvia atricapilla*) that undertakes biannual migratory night flights of up to 6000 km between Europe and Africa using the Earth’s magnetic field (Viehmann 1979; Schwarze *et al*. 2016; Kobylkov *et al*. 2020; Kobylkov *et al*. 2022; Haase *et al*. 2022; Leberecht *et al*. 2022; Leberecht *et al*. 2023). Recently, studies on the neural mechanisms underlying magnetoreception in night-migratory birds have led to the identification of magneto-processing areas in the visual and trigeminal systems (Heyers *et al*. 2007; Heyers *et al*. 2010; Heyers *et al*. 2022; Zapka *et al*. 2009; Zapka *et al*. 2010; Hein *et al*. 2010; Lefeldt *et al*. 2014; Mouritsen and Ritz 2005; Mouritsen, Heyers, and Güntürkün 2016; Mouritsen 2018; Elbers *et al*. 2017; Günther *et al*. 2018; Günther *et al*. 2021; Günther *et al*. 2024; Kobylkov *et al*. 2020; Haase *et al*. 2022) as well as higher brain areas that potentially integrate multimodal navigational information (Kobylkov *et al*. 2022; Heyers *et al*. 2022). To date, however, scientists lack a standardized brain reference atlas for comparative neuroanatomical studies on migratory bird species, which could enhance our understanding of the brain regions and mechanisms involved in neuronal processing of navigational cues.

Historically, 2D print-based atlases have been used to identify brain areas across different animal species (Wullimann, Rupp, and Reichert 1996; Paxinos 2000; Paxinos and Franklin 2001; Swanson 2004; Puelles *et al*. 2019). However, print-based atlases hinder the reliable and reproducible mapping of brain areas, since they are digitally inaccessible and do not allow the alignment of data to a common reference. Furthermore, the constant discovery of new and/or functionally defined brain areas and the subdivision of existing ones quickly render print-based atlases outdated. In contrast, using automatic sample-to-atlas registration creates unprecedented opportunities for integrating data from multiple brains into a unified reference system. Therefore, there is a critical need for 3D digital brain atlases that provide a standardized coordinate framework for the reliable definition of brain areas (Papp *et al*. 2014; Kim *et al*. 2015; Wang *et al*. 2020; Kenney *et al*. 2021; Tyson and Margrie 2022; Montague *et al*. 2023; Shainer *et al*. 2023; Gustison *et al*. 2024). The generation of high resolution digital reference atlases to systematically map brain areas (both anatomically and functionally) as well as their open-source availability, has dramatically facilitated reproducibility of results across laboratories. This has significantly advanced systems neuroscience research in all animal models for which these types of atlases have been developed. Moreover, modern experimental neuroscience relies on techniques such as device implantation and specific cellular marker expression for the stimulation and recording of neurons (Adelsberger *et al*. 2014; Steinmetz *et al*. 2021). The precise localization of viral injections or implanted devices within the brain is therefore critical to validate the outcome of such experiments (Tyson *et al*. 2022). Additionally, an increasing body of research relies on the precise localization and registration of marker expression and connectivity in a common reference framework (Tyson *et al*. 2021; Tyson and Margrie 2022).

Digital, interactive 3D atlases already exist for common model-organisms such as the mouse (Wang *et al*. 2020) or zebrafish (Kenney *et al*. 2021). These atlases, along with many others, have been incorporated into the BrainGlobe Atlas API (Claudi *et al*. 2020) enabling automatic registration of sample brains or mapping implanted devices and labelled cell populations in a common reference space using open-source processing tools (Niedworok *et al*. 2016; Tyson *et al*. 2021; Tyson *et al*. 2022). For bird brains, all presently available online atlases have significant limitations. 2D photo-based atlases for the zebra finch, derived from gene expression profiles or histological stainings, lack the possibility to visualize more than two orientations in the same brain (Nixdorf-Bergweiler *et al*. 2007; Karten *et al*. 2013; Lovell *et al*. 2020). In addition, low spatial resolution 3D atlases have been developed for the zebra finch (Poirier *et al*. 2008) (80 × 160 × 160 *µm*voxel size), the pigeon (Güntürkün *et al*. 2013) (60 × 60 × 60 - 222 × 223 × 223 *µm* voxel size) the starling (De Groof *et al*. 2016) (223 × 223 × 223 *µm* voxel size), the canary (Vellema *et al*. 2011) (40 × 40 × 40 - 60 × 60 × 60 *µm* voxel size) and the Japanese quail (Yebga Hot *et al*. 2022) (150 × 150 × 150 *µm* voxel size) using magnetic resonance imaging (MRI). Importantly, however, these resources either have restricted access, are based on a single specimen and/or lack high spatial resolution.

To overcome the resulting methodological limitations, we have developed the first digital, histology-based, 3D brain atlas for a bird species, the migratory Eurasian blackcap, using data from multiple specimens, at an isotropic voxel size of 25 *µm*. Here, we describe the entire process involved in generating this atlas, from whole-brain data acquisition using serial two-photon tomography, to the creation of an average template and expert neuroanatomical segmentation. In total, we segmented 24 brain areas from which 19 are commonly found across all bird species. Given the magneto-sensing capabilities of the Eurasian blackcap, we additionally segmented 5 magneto-processing brain areas. We also incorporate the Eurasian blackcap atlas into the BrainGlobe Atlas API, thus making it “ready-to-use” within the BrainGlobe ecosystem. This integration provides an unified data analysis ecosystem for tasks such as cell and marker detection (Tyson *et al*. 2021), registration of experimental data to the atlas (Tyson *et al*. 2022) as well as data visualization and exploration (Claudi *et al*. 2021).

## 2 Results

### General approach

The overall strategy and steps we performed to create the backcap atlas are outlined in Figure 1. First, we employed serial two-photon (STP) tomography to image and cut whole bird brains. Next, the individual 3D images from different brains were iteratively aligned and averaged to create a representative brain template. Following this, manual expert segmentation and annotations were performed on the generated average 3D template. Finally, the atlas was incorporated into the BrainGlobe Atlas API ecosystem and automatic registration, cell detection as well as object mapping were demonstrated on experimental data.

**Figure 1.**
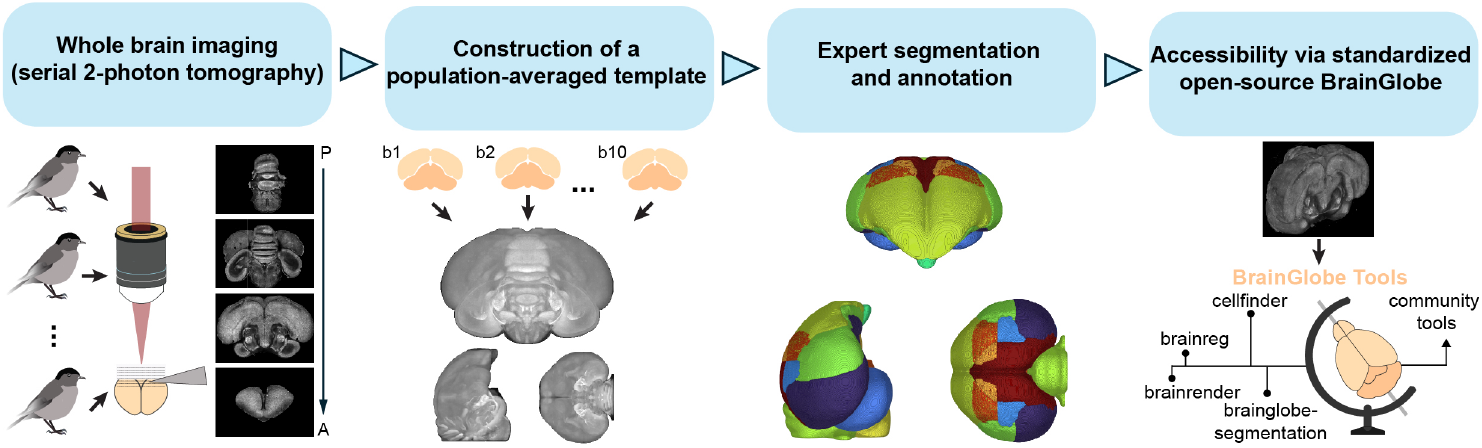
Overview of the approach for creating a 3D bird brain atlas for the night-migratory Eurasian blackcap (*Sylvia atricapilla*). Acquisition of high resolution images of entire perfused brains (A: Anterior, P: Posterior) using automated serial two-photon tomography. Construction of an average template brain using the individual brains (b1-b10). Hierarchical segmentation of volumes into neuroanatomical brain compartments. The resulting 3D bird brain atlas is made accessible via the BrainGlobe ecosystem, enabling, for example, automated cell detection and registration of experimental data to the atlas.

### Serial section two-photon whole brain imaging

Ten wild-caught adult blackaps were transcardially perfused and their brains harvested for automated STP tomography, which is a two-photon microscope equipped with a vibrating blade microtome (Figure 2A,B). This method produced well-aligned, high-resolution images (2 × 2 × 5 *µm* voxel size) of entire bird brains (Figure 2C-F), including labelled neurons in five out of the ten specimens where small viral injections (∼100 nl) into the brain had been performed.

**Figure 2.**
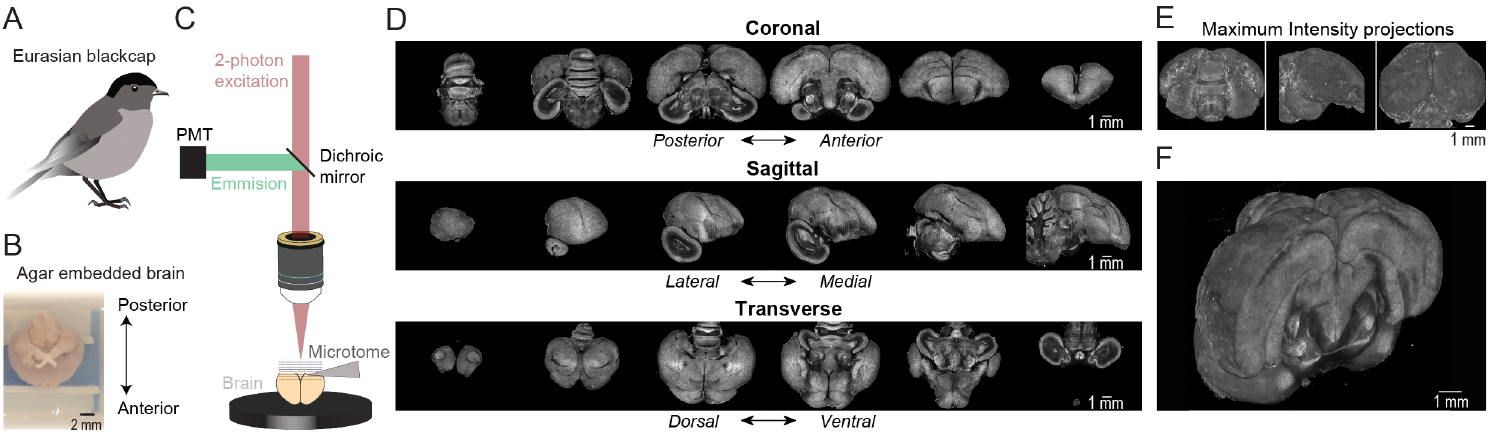
Imaging of whole bird brains using serial two-photon tomography. (A) Schematic of the Eurasian black cap (*Sylvia atricapilla*). (B) Brain embedded in 5% agar with ventral side facing upwards. (C) Simplified schematic of automated serial two-photon microscope used for whole-brain imaging. (D) Coronal (top), sagittal (middle) and transverse (bottom) slice views of an individual brain. (E) Coronal, sagittal and transverse view of the 3D volume generated for an individual brain visualized using a maximum intensity projection. (F) 3D volume of a single bird brain visualized using a maximum intensity projection. Note that the brain is only partially displayed for illustration.

### Construction of a population-averaged brain template

Our goal was to create an average brain template that represents the population as a whole, without bias toward the unique features of any single individual. The challenges in creating an average stem from individual variations in brain shape and size, as well as offsets in the imaging field of view. Therefore, the images from individual animals have to be registered into the same target space and then averaged. However, the choice of the target space may introduce undesired bias: for example, if all brains are aligned to a single target brain, the resulting average will acquire the shape of the target. To overcome this limitation, we used the symmetric group-wise normalization (SyGN) algorithm (Avants *et al*. 2011; Fonov *et al*. 2011), a method commonly employed in MRI to generate high-quality, unbiased average templates for both human (Avants *et al*. 2010) and animal brains (Love *et al*. 2016; Seidlitz *et al*. 2018; Jung *et al*. 2021; Liu *et al*. 2021).

The SyGN algorithm iteratively registers and averages multiple individual brain images to a common target space (see Supplementary Figure 1 for a schematic). It begins with an initial template image - i.e. one of the sample brains - which is refined through successive iterations in both intensity and shape, ultimately converging on an unbiased average representation (Supplementary Figure 2). Specifically, the algorithm proceeds in five main steps: (1) each individual specimen is registered to the starting template, (2) the images are transformed into the current template space, (3) the voxel-wise average of the transformed images is calculated and the resulting average intensity image is sharpened, (4) the individual-to-template transformations are averaged and then inverted, yielding an average template-to-individual transformation, and (5) this inverted transformation is applied to the sharpened average from step 3, updating the template’s shape to better reflect the group average. The updated template then becomes the new registration target for the next iteration. Repeating step 5 over multiple iterations ensures that the final average is no longer biased toward the starting template. Moreover, the final template benefits from the increased signal-to-noise ratio gained through averaging (thanks to step 3) while avoiding blurring, thereby preserving fine anatomical details (Avants *et al*. 2010).

Here we constructed a brain template using the ten blackcap brain image stacks acquired via STP tomography. The images were prepared by cropping, downsampling to 25 *µm* isotropic resolution, and correcting for intensity inhomogeneities. To maximize our data set, we divided each brain into left and right hemispheres and symmetric images were created by mirroring each hemisphere. After excluding two damaged hemispheres, 18 left-right symmetric brain images were used as inputs for the SyGN algorithm (Supplementary Figure 2A). This approach ensured left-right symmetry in the resulting template, which is crucial for certain applications. For instance, asymmetric templates cannot be used to estimate left-right anatomical differences in a population, as any observed differences may be confounded by the inherent asymmetry of the template itself (Fonov *et al*. 2011). Further details on image preparation and template construction using SyGN can be found in the Methods section.

The final template generated by this process is shown in Figure 3A and, juxtaposed with individual images that contributed to its construction. When comparing the averaged template to single specimens we found that individual anatomical landmarks and borders between brain areas were much sharper (Figure 3). Moreover, peculiarities of individual brains such as injection tracks and labelled cells were absent from the averaged template (Figure 3B). Taken together, the use of the SyGN algorithm produces an artefact-free and high-contrast average brain template serving as an ideal registration target and annotation basis.

**Figure 3.**
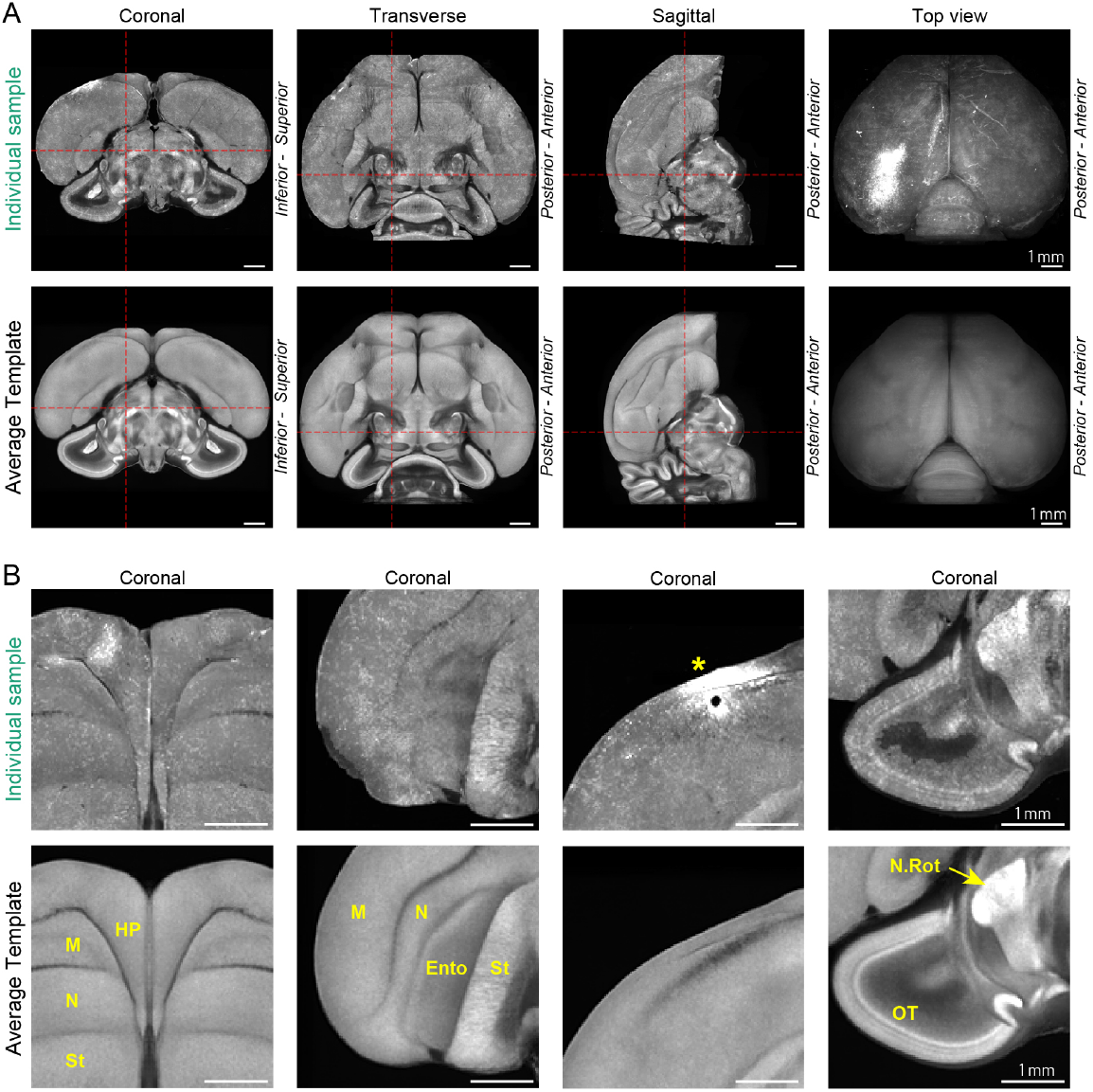
Average brain template. (A) Individual example brain (top) is juxtaposed with the average template (bottom) constructed from the ten brains (18 hemispheres). Three orthogonal slices—coronal, transverse, sagittal—are shown alongside a top view of the brain (maximum intensity projection, attenuated with an exponential factor of 0.01 to emphasize features closest to the top surface). (B) A closer look at selected coronal slice views of the example brain (top) and the average template (bottom). Borders between anatomical subdivisions are much sharper on the average template. Moreover, the average brain is free of the tissue damage visible on the example brain—see the location of an anatomical tracer injection into the example brain (asterisk). M: Mesopallium; N: Nidopallium; St: Striatum; HP: Hippocampus; Ento: Entopallium; N.Rot: Nucleus Rotundus; OT: Optic Tectum.

### Expert segmentation and annotation

The generated average template (Figure 4A) was used to annotate six main brain compartments (Cerebellum (Cb), Diencephalon (Di), Mesencephalon (Me), Pallium (P), Pons (Pn) and Striatum (St)). The delineation of these brain areas was based on anatomical landmarks with the help of 2D images of Nissl- and myelin-stained brain sections from the zebra finch atlas by Karten et al. (Karten *et al*. 2013), a phylogenetically related songbird species to the blackcap. The annotations were thereby performed on a single hemisphere. These main brain areas were then further subdivided into conspicuous brain areas common to all bird species for which atlases exist, i.e. the Nucleus Rotundus (N. Rot.) in the diencephalon, the Optic Tectum (OT) in the mesencephalon and the Arcopallium (A), Dorsolateral Corticoid Area (CDL), Entopallium (Ento), Hyperpallium (H), Hippocampus (HP), Mesopallium (M), Nidopallium (N) and Olfactory Bulb (OB) in the pallium. In addition to these anatomically defined brain areas, we annotated five main functionally defined brain areas involved in magnetic field processing based on previous studies (Günther *et al*. 2018; Günther *et al*. 2021; Günther *et al*. 2024; Mouritsen and Ritz 2005; Mouritsen, Heyers, and Güntürkün 2016; Mouritsen 2018) (Figure 4D). These included “Cluster N” (in the following subdivided into a hyperpallial and a hippocampal part) (Heyers *et al*. 2007; Heyers *et al*. 2022; Zapka *et al*. 2009; Zapka *et al*. 2010; Hein *et al*. 2010), the diencephalic Nucleus geniculatus lateralis pars dorsalis (Gld) (Heyers *et al*. 2007), the dorsal and ventral principal sensory trigeminal brainstem nuclei (PrVd, PrVv) (Heyers *et al*. 2010; Lefeldt *et al*. 2014; Elbers *et al*. 2017; Haase *et al*. 2022), the lateral and medial spinal sensory trigeminal brainstem nuclei (SpVl, SpVm) in the pons (Heyers *et al*. 2010; Lefeldt *et al*. 2014; Elbers *et al*. 2017; Haase *et al*. 2022) and the telencephalic frontal nidopallium (NFT) (Kobylkov *et al*. 2020). Our segmentation resulted in a 3D model of the brain (Figure 4C). Each brain area was given a unique abbreviation and color associated with an anatomical hierarchy (Figure 4B). The location and boundaries of these magneto-responsive areas were ultimately determined by an expert in the magnetoreception field (see methods) using either expression of immediate early genes (Egr-1; (Jarvis and Nottebohm 1997)) that serve as proxies for neuronal activation during orientation/magnetic field stimulation or neuronal tracing data (Heyers *et al*. 2007; Heyers *et al*. 2010; Heyers *et al*. 2022; Elbers *et al*. 2017; Kobylkov *et al*. 2020).

**Figure 4.**
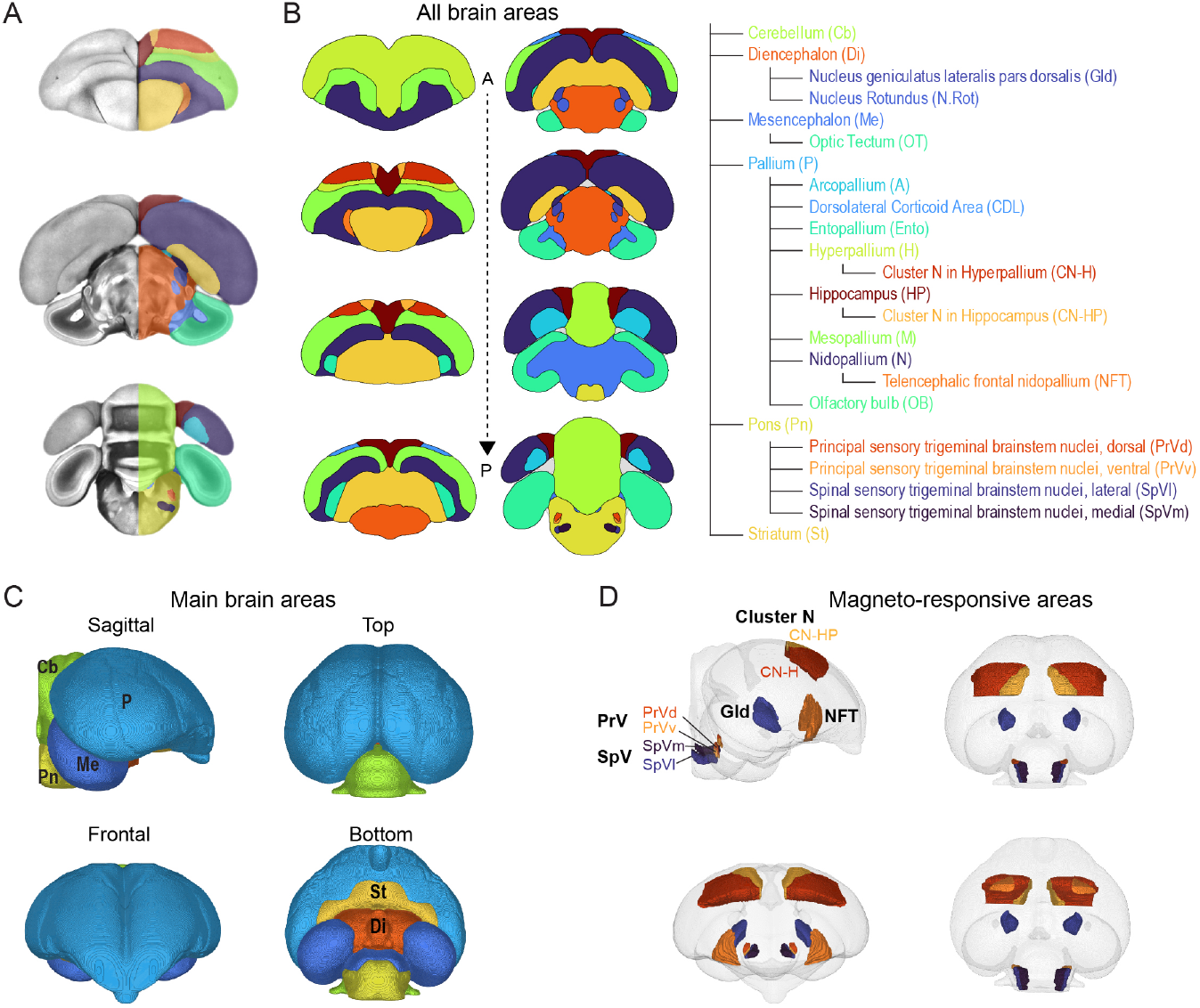
Manual segmentation and annotation of brain areas. (A) Coronal slice views of the average template brain overlaid with manual segmentation of brain areas on one hemisphere. (B) Coronal sections with individual segmented brain areas along the anterior-posterior axis (left). Hierarchical tree structure of the brain segmentation (right), showing color-coded brain area names and their abbreviations. (C) 3D rendering of the brain with the six main brain subdivisions based on anatomical landmarks. (D) 3D rendering of the brain with segmented magneto-responsive areas. Note that these brain areas are subdivided based on anatomical and functional landmarks.

### Online accessibility and BrainGlobe toolbox compatibility

The finalized atlas is freely available at https://gin.g-node.org/BrainGlobe/atlases and can be used programmatically through the BrainGlobe Atlas API (Claudi *et al*. 2020) under the atlas name “eurasian_blackcap_25_ *µm*”. The atlas is available for use within the BrainGlobe ecosystem (https://brainglobe.info/documentation) and other open-source tools built by the community (https://brainglobe.info/community/externaltools) such as ABBA (Chiaruttini et al. 2024). The atlas is a live resource and updates as well as tutorials will be available at https://brainglobe.info/blackcap.

To demonstrate the usability of our atlas within the BrainGlobe ecosystem, we conducted automatic registration, cell detection, and data visualization using a sample brain injected with a retrograde viral tracer (scAAV-DJ/9/2-EGFP) in the magneto-responsive brain area, Cluster N (Figure 5). We successfully registered the sample brain to our reference atlas and used BrainGlobe’s cellfinder tool to automatically detect and quantify cells (Figure 5A,B). Additionally, we manually segmented the injection track within the sample brain using the BrainGlobe tool brainglobe-segmentation. Finally, we rendered and visualized all data registered to the reference atlas using the brainrender software (Figure 5C). We identified multiple brain areas projecting to Cluster N (Figure 5D) and quantified the number of cells within these regions, which largely corroborated previously demonstrated telencephalic connectivities of Cluster N (Heyers *et al*. 2007; Heyers *et al*. 2022). We thereby demonstrate the usability and applicability of our atlas within the BrainGlobe ecosystem (Figure 5D). Taken together, we successfully implemented and tested our blackcap brain atlas in the BrainGlobe ecosystem which will allow users to compare their own data across individual brains, experimental modalities and laboratories.

**Figure 5.**
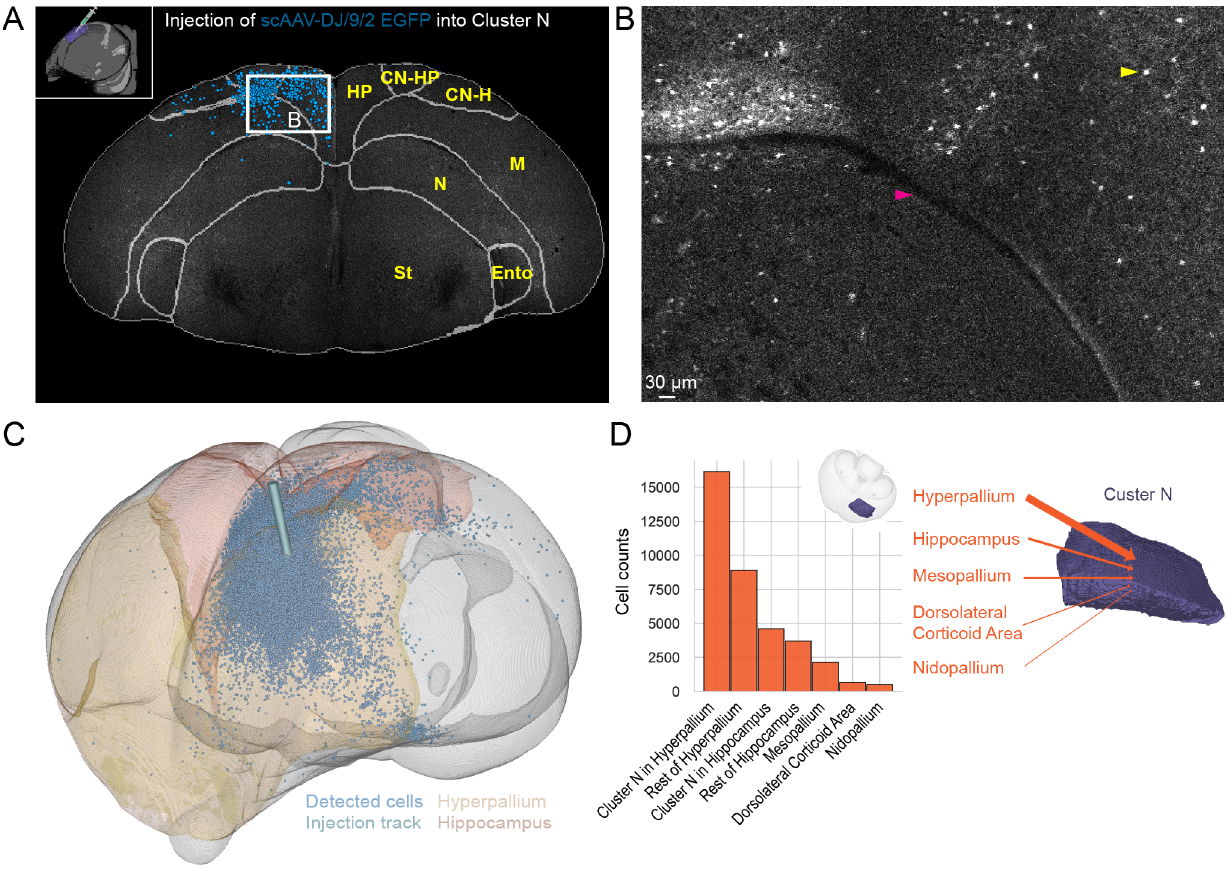
Application of the BrainGlobe toolbox for brain registration, automated cell detection and object segmentation. (A) Schematic depicting the viral injection of scAAV-DJ/9/2 EGFP into the magneto-responsive area Cluster N (top left insert). Automatically detected cells overlaid on raw data along with the brain region segmentation. HP: Hippocampus; CN-HP: Cluster N in Hippocampus; CN-H: Cluster N in Hyperpallium; M: Mesopallium; N: Nidopallium; St: Striatum; Ento: Entopallium. (B) Enlarged insert of brain area from A displaying labelled cells and anatomical features. Example cell highlighted with yellow arrowhead. Border between Hippocampus and Mesopallium is highlighted with a magenta arrowhead. (C) Visualization of detected cells warped to the bird brain atlas coordinate space in 3D along with the injection track and two brain areas (Hyperpallium and Hippocampus). (D) Bar plot displaying total number of cells detected per brain area (left) and schematic illustrating main projections onto Cluster N (right). Thickness of the arrows indicates projection strength.

## 3 Discussion

Here we provide the neuroscience research community with the first digital, high-resolution 3D bird brain atlas based on light microscopy that enables the precise delineation of brain structures.

This resource was made possible by leveraging automated STP tomography (Ragan *et al*. 2012; Osten and Margrie 2013), enabling near-perfect alignment between consecutive brain slices since the imaging is carried out on the specimen block face prior to the sections being cut. STP tomography also eliminates the need for tissue clearing prior to imaging, thus avoiding significant tissue shrinkage. As a result, the imaging quality across individual brains remains consistently high, with minimal variation and photobleaching, a crucial factor when working with animal species where only a limited number of brain samples are available for atlas construction, such as night-migratory passerines that have proven difficult to breed in captivity.

To create an average brain template, we combined male and female bird brains of similar age and ensured symmetry by mirroring each hemisphere along the mid-sagittal plane. Given the limited availability of migratory bird species, which need to be wild-caught, the symmetric, all-age, all-gender template constructed here serves as a valuable starting point. This bears the potential caveat that e.g. known brain anatomical sexual dimorphisms, such as in brain areas involved in vocal control, (e.g. HVC and/or Areas X in the songbird brain, Nottebohm and Arnold 1976), are averaged out in the template. However, several known structural lateralizations in birds are constituted by left-right differences of axonal pathways such that lateralized functions emerge from systems interactions (Güntürkün *et al*. 1998; Manns and Ströckens 2014; Letzner, Manns, and Güntürkün 2020) as well as cellular coding properties (Xiao and Güntürkün 2018; Costalunga, Vallentin, and Benichov 2024) and are therefore usually not reflected in volumetric differences (Güntürkün 1997; Güntürkün, Ströckens, and Ocklenburg 2020).

The Eurasian blackcap brain atlas developed here represents the first-ever comprehensive atlas for a migratory bird species, making it an indispensable resource for researchers studying the neuroanatomical and functional neuronal mechanisms of long-distance bird migration, orientation and navigation as well as magnetic field sensing (Mouritsen 2018; Nimpf and Keays 2022). Beyond its critical relevance to migration, orientation, navigation and magnetic sensing, this atlas also offers substantial advantages compared to any previously available whole-brain bird atlases and will help the wider neuroscience community working with other bird species as a model organism. In contrast to other digitally available bird atlases such as the zebra finch histological brain atlas (Karten *et al*. 2013), where different brains were sectioned in either the sagittal, transverse, or horizontal plane, our atlas is based on whole-brain 3D imaging, which allows visualization from any arbitrary perspective within a common coordinate space. Additionally, our current version of the atlas is based on the average of ten bird brains, which allows a much more precise anatomical delineation as compared to single specimens. Our atlas also provides a spatial voxel resolution of 25 *µm*, a significant improvement over the resolution of any 3D digital bird atlas currently available online (Poirier *et al*. 2008; Güntürkün *et al*. 2013; De Groof *et al*. 2016). At the same time, the current spatial resolution remains compatible with standard computer hardware, allowing users to efficiently register their own data. Lastly, our atlas features annotations of magneto-responsive regions, thus expanding our understanding of these functionally defined avian brain areas.

The digital and open-source nature of the Eurasian blackcap brain atlas developed here allows for wider accessibility and utility across a broader spectrum of research fields, including comparative neuroanatomy, behavioral neuroscience, and evolutionary biology. In the first version of the atlas, we segmented 24 brain areas with borders that could be delineated by a small number of human annotators. Given that manual annotation is subjective, unreliable (Niedworok *et al*. 2016) and time- and labor-intensive, making the data and atlas accessible to a larger number of expert bird neuroanatomists will help distribute the workload and facilitate consensus on the characterization of specific brain areas. Most importantly, advances in whole brain antibody labelling and staining procedures, paired with serial STP tomography, will further guide brain area subdivision based on region-specific expression of marker genes or proteins as has been performed in other animal species (Kenney *et al*. 2021). Finally, recent developments in supervised machine learning methods (Essen and Glasser 2018) hold promise for automated delineation of brain areas in the future (Iqbal, Khan, and Karayannis 2019; Billot *et al*. 2023). Importantly, this atlas will be regularly updated to incorporate any new data.

This atlas, made possible through advanced imaging techniques, sets a new benchmark in the quality and consistency of avian brain atlases, ensuring that researchers can rely on it as a reference point. The methodology used here will serve as a comprehensive blueprint for the development of similar atlases for other animal species in the future.

## 4 Methods

### Animals and housing

All animal procedures were approved by the local and national authorities for the use of animals in research (Niedersächsisches Landesamt für Verbraucherschutz und Lebensmittelsicherheit/LAVES, Oldenburg, Germany, Az.: 33.19-42502-04-20/3492; 33.19-42502-04-21/3650). Ten adult Eurasian blackcaps (*Sylvia atricapilla*) of both sexes (two females and eight males, greater than one year old) were caught by mist-netting in the vicinity of Oldenburg University. Birds were then housed in pairs and experienced a circadian and circannual light regime matching the local natural conditions. Food and water were provided ad libitum. To obtain the most accurate average brain dataset possible given the limited sample size of wild-caught specimens, the brains of male and female individuals were pooled. This does not mean to imply that the brain of *Sylvia atricapilla* is not sexually dimorphic and that further development of individual male and female atlases could not prove useful. From these ten brains, five had received viral injections and therefore displayed minor tissue damage along the injection track and contained some labelled neurons. Due to extensive damage around the viral injection site (Supplementary Figure 2A) two out of twenty hemispheres (1 male, 1 female) were excluded.

### Perfusion and tissue extraction

Eurasian blackcaps were deeply anaesthetized with pentobarbital (Narcoren, Boehringer Ingelheim, Ingelheim, Germany) and transcardially perfused with 0.9% saline followed by paraformaldehyde (4% in phosphate buffered saline (PBS), pH 7.4). Brains were extracted from the skull and stored in 100 mM PBS at 4 °C until being ready for imaging.

### Brain wide serial two-photon imaging

For serial section two-photon imaging, on the day of imaging, brains were removed from the PBS and dried. Brains were then embedded in agarose (5%) using a custom alignment mould to ensure that the brain was perpendicular to the imaging axis. The agarose blocks containing the brains were trimmed and then mounted onto the serial two-photon microscope containing an integrated vibrating microtome and motorized x–y–z stage (STP tomography, Ragan *et al*. 2012; Osten and Margrie 2013). For this, a custom system controlled by ScanImage (v5.6, Vidrio Technologies, USA) using BakingTray (Campbell 2020) was used. Imaging was performed using 920 nm excitation illumination and fluorescence emission was band-pass filtered with either 640-570 nm for red, 550-505 nm for green and 491-437 nm for blue. Images were acquired with a 2.3 × 2.3 *µm* pixel size, and 5 *µm* plane spacing. Ten optical planes were acquired using a piezoelectric motor over a depth of 50 *µm* in total. To image the entire brain, images were acquired as tiles and stitched using StitchIt (Campbell, Blot, and lguerard 2020). After each mosaic tile was imaged at all ten optical planes, the microtome automatically cut a 50 *µm* slice, enabling imaging of the subsequent portions of the sample and resulted in full 3D imaging of entire brains. The images in the blue, red and green channels were saved as a series of 2D TIFF files.

### Preparation of images for template construction

We used the set of ten blackcap brain image stacks obtained through STP tomography. We visually inspected the data to select channels with a good signal-to-noise ratio and clear contrast between gray and white matter structures. In 8/10 cases, we used the autofluorescence emitted at 505 – 550 nm, while in the remaining 2/10 cases, we used the autofluorescence emitted at 437 – 497 nm.

We manually cropped each image to tightly enclose the brain tissue and downsampled them to an isotropic resolution of 25 *µm*. The images were reoriented to conform to BrainGlobe’s ASR convention—placing the origin at the anterior superior right corner and ordering the axes as anterior-posterior, superior-inferior, and right-left (Claudi *et al*. 2020). The reoriented images were then converted from TIFF to NIfTI format for compatibility with the Advanced Normalization Tools (ANTs) software suite (Tustison *et al*. 2021). To correct for intensity inhomogeneities, ANTs’ N4 bias field correction function was applied (Tustison *et al*. 2010). Brain masks were generated using scikit-image (Walt *et al*. 2014). The images were blurred with a Gaussian kernel (standard deviation of 150 *µm*) and thresholded using the ‘triangle’ method. The largest connected component was selected from the thresholded mask, and binary closing was applied with a square footprint of 250 *µm* to eliminate small holes.

To ensure symmetry and exclude two damaged hemispheres, we virtually hemisected each brain image along the mid-sagittal plane. This was achieved by:

- Defining the mid-sagittal plane in one image by annotating multiple midline points with napari (Ahlers et al. 2023) and fitting a 2D plane.
- Rotating the image in 3D so its mid-sagittal plane aligned with the central sagittal plane of the image volume.
- Rigidly aligning each individual image to this rotated target using ANTs registration, ensuring consistent alignment of the mid-sagittal plane across subjects.
- Splitting aligned images into left and right hemispheres along the mid-sagittal plane, and discarding the two damaged hemispheres.
- Mirroring the 18 intact hemispheres along the left-right axis to generate symmetric images.

These steps produced 18 symmetric brain images and their corresponding masks, ready for template construction. The code used for image preparation has been incorporated into the *brainglobe-template-builder* Python package (Sirmpilatze et al. 2025).

### Iterative template construction with SyGN

The symmetric brain images and their masks served as inputs to the symmetric group-wise normalization (SyGN) algorithm for unbiased average template construction (Avants *et al*. 2011). The brain masks were employed during registration to focus computations within brain areas and exclude background.

SyGN is canonically implemented in the open-source toolkit Advanced Normalization Tools (ANTs) (Tustison *et al*. 2021) through the *antsMultivariateTemplateConstruction2*.*sh* script. Here we used ANTs v2.5.2 in conjunction with an optimized implementation of the template construction script (*optimized_antsMultivariateTemplateConstruction v1*.*0*), which offers enhanced features such as the ability to resume interrupted computations and automatic adjustment of registration parameters based on input image size and resolution (Devenyi 2024). Template construction was initialized using one of the input images as a starting template and executed in four stages with progressively higher-order transformation types: rigid (6 parameters), similarity (6 rigid parameters plus 1 uniform scaling), affine (12 parameters), and non-linear. The non-linear stage relied on ANTs’ symmetric diffeomorphic normalization (Avants *et al*. 2008), which is among the top-performing algorithms for topology-preserving non-linear image registration (Klein *et al*. 2009). Each stage was passed through four iterations of the SyGN algorithm. Briefly, each SyGN iteration involved the following steps (see schematic in Supplementary Figure 1):

1. Register each individual subject to the current template using linear and/or non-linear transformations (depending on the stage).
2. Transform the individual images to the current template space using the computed matrices and/or nonlinear deformation fields.
3. Calculate the voxel-wise trimean of the transformed images and apply a sharpening filter to the resulting average intensity image.
4. Average the individual-to-template transformations from step 1 to obtain an average transformation, then invert it.
5. Apply the inverted average transformation from step 4 to the sharpened average intensity image from step 3. This step updates the template’s shape to more closely reflect the average shape of the individual brains. The updated template becomes the new registration target for the next iteration.

The average images from each stage and iteration were visually inspected to ensure convergence to a stable template (see Supplementary Figure 2). The final template, generated after the 4th iteration of the non-linear stage, was selected as the reference image for the blackcap brain atlas. Note that whereas the original SyGN implementation computes the average intensity image by taking the voxel-wise mean of the transformed images, we instead used the voxel-wise efficient trimean—a weighted average of the 20th, 50th (median) and 80th percentiles: (P20 + 2P50 + P80) / 4. This makes the average more robust to outliers, i.e. less affected by exceptionally bright or dark pixels that may be present in any given brain image. We found this approach to be especially beneficial in our case, to prevent injection tracks and labelled cells (which were present in some samples) from contaminating the average image.

### Segmentation

The segmentation and annotation of brain areas was manually performed by expert bird brain anatomists using ITK-SNAP http://www.itksnap.org/pmwiki/pmwiki.php, a freely available software package for working with multimodal images. The annotations were done on a single hemisphere which was subsequently reflected to ensure a fully symmetric annotation image. The annotations were performed by members of the Animal Navigation group at the University of Oldenburg. All annotations were ultimately checked by Dominik Heyers, an expert in avian brain anatomy for two decades.

### Atlas packaging

The final template and annotation images were packaged into the BrainGlobe format. Briefly, the metadata from ITK-SNAP was converted to a standardized format (Claudi *et al*. 2020). A 3D mesh was then generated for each segmented brain region using SciPy (Virtanen *et al*. 2020), PyMCubes (https://github.com/pmneila/PyMCubes) and Vedo (Musy et al. 2022). The packaging script can be found at https://github.com/brainglobe/brainglobe-atlasapi.

### Neuronal Tracing

For neuronal tracing, birds were fully anesthetized using IsofluraneCP© (cppharma, Burgdorf, Germany) at a concentration of 1–1.5Vol.% administered through a beak mask and head-fixed in a custom-built stereotactic apparatus. The surfaces of both telencephalic hemispheres and the cerebellum were positioned in the same horizontal plane resulting in an angle of the plane between the tip of the beak and the ear bars of approximately 45º below the horizontal zero plane of the apparatus. The birds’ scalp was anesthetized using a surface anesthetic (Xylocain; Astra Zeneca, Wedel, Germany), incised, and temporarily pulled aside. Tracer injections into the center of Cluster N were achieved by using coordinates (anterior/posterior (x): 5.0 mm; medio-lateral (y): 3.0 mm; depth (z): 0.5 mm) of the target structure relative to the confluence of the superior sagittal and cerebellar “Y” blood sinus providing the zero coordinate. Viral tracer (scAAV-DJ/9/2 EGFP; ETH Zurich, VVF Cat#v65) was administered through a small window in the skull above Cluster N by minute pressure injections of 200-500 nl using a microinjector (WPI-2000, World Precision Instruments, USA) and bevelled glass capillaries. After the surgery, the skull and skin were repositioned and sealed with cyanoacrylate surgical glue (Histoacryl®, BRAUN, Rubi, Spain). Post-surgical analgesia was provided through intramuscular administration of Metacam© (0.1 ml/kg body weight in 0.9% NaCl, Boehringer Ingelheim, Ingelheim, Germany) for 72 h. Each bird was given 3 weeks to allow for viral expression.

### Tracing data analysis and visualization

Following imaging, the brain was registered to the atlas using brainreg (Tyson *et al*. 2022). All parameters were the defaults other than Gaussian smoothing which was set to 0 and the number of histogram bins used for the calculation of Normalised Mutual Information which was set to 64 for both the sample image and the atlas template. Labelled presynaptic cells were detected using cellfinder (Tyson *et al*. 2021) using all default settings and the default classification model. Detected cells were manually corrected using napari (Ahlers et al. 2023). The coordinate of the injection site was determined using brainglobe-segmentation (Tyson *et al*. 2022), and this was visualized alongside the detected cells in the atlas using brainrender (Claudi *et al*. 2021).

## 5 Author Contributions

S.W. conceived the project with input from A.L.T., N.S., A.F., D.H. and H.M.. S.W. imaged all brains and generated 3D volumes. N.S. and A.F. performed the image processing and the average template construction. D.H. performed perfusions and D.A., L.S., K.H., I.M. and D.H. annotated brain areas. A.F. and A.L.T. integrated the atlas into the BrainGlobe ecosystem of tools. A.L.T. performed data analysis. All authors significantly contributed to manuscript writing.

## Acknowledgments

The authors are grateful to the members of the Advanced Microscopy Facility at the Sainsbury Wellcome Centre and to Mateo Velez-Fort for comments on the manuscript. S.W. was funded by a Feodor-Lynen fellowship from the Alexander von Humboldt Foundation. S.W. was also funded by Wellcome (219627/Z/19/Z; 214333/Z/18/Z) and Gatsby Charitable Foundation, GAT3755) awarded to T.W.M.. A.L.T. and N.S. were funded by the core grant to the Sainsbury Wellcome Centre (Wellcome - 219627/Z/19/Z, Gatsby Charitable Foundation - GAT3755) and A.L.T. was funded by the core grant to the Gatsby Computational Neuroscience Unit (Gatsby Charitable Foundation - GAT3850). A.F. was funded by grant 2022-309537 from the Chan Zuckerberg Initiative DAF, an advised fund of Silicon Valley Community Foundation. D.H. and H.M. received valuable financial support by Deutsche Forschungsgemeinschaft (SFB 1372 “Magnetoreception and Navigation in Vertebrates”, project number 395940726 to D.H. and H.M., employing D.A., L.S., K.H. and I.M.) and European Research Council (under the European Union’s Horizon 2020 research and innovation program, grant agreement no. 810002 (Synergy Grant: “QuantumBirds”) to H.M..

## Data, Materials, and Software Availability

Code to create the brain template can be found at https://github.com/brainglobe/brainglobe-template-builder and code to package the final atlas can be found at https://github.com/brainglobe/brainglobe-atlasapi. Raw data are available on request.

