## Supporting information for "Mapping the magnetoreceptive brain: A 3D digital atlas of the migratory bird Eurasian blackcap (*Sylvia atricapilla*)"

### A digital three-dimensional brain atlas of a magneto-receptive migratory bird, the Eurasian blackcap (*Sylvia atricapilla*)

Nikoloz Sirmpilatze, Alessandro Felder, Dinora Abdulazhanova, Leonard Schwigon, Katrin Haase, Isabelle Musielak, Troy W. Margrie, Henrik Mouritsen, Dominik Heyers, Adam L. Tyson, Simon Weiler

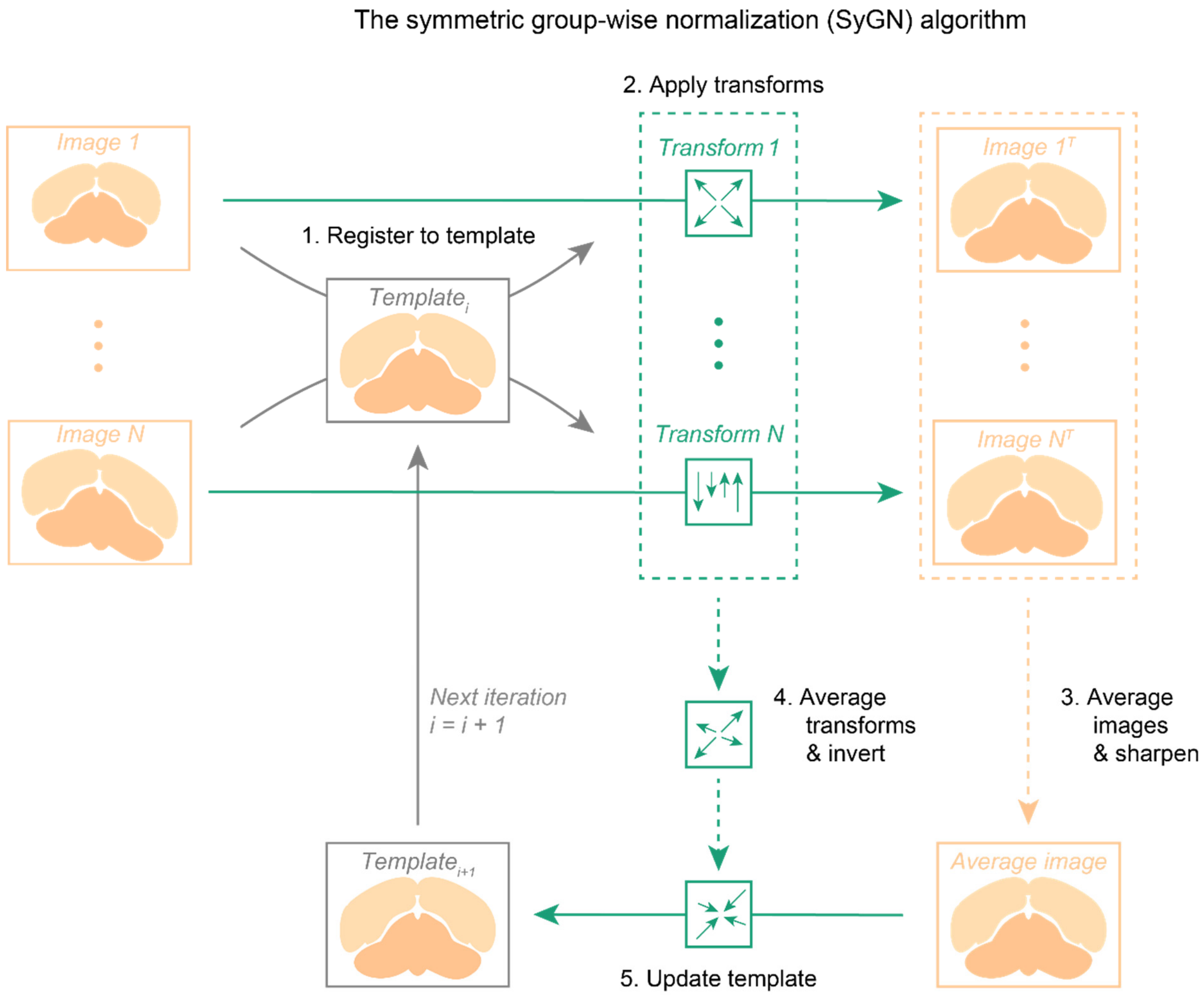


**Supplementary Figure 1: The symmetric group-wise normalization (SyGN) algorithm.** The schematic depicts a single iteration of the SyGN algorithm for unbiased average template construction (Avants et al., 2011). **(1)** Each individual brain image is registered to the current template using linear and/or non-linear transformation (depending on the transform stage, see Supplementary Figure 2). **(2)** The individual images are transformed to the current template space using the computed matrices and/or nonlinear deformation fields. **(3)** The transformed images are averaged voxel-wise and a sharpening filter is applied to the resulting average intensity image. **(4)** The individual-to-template transformations from step 1 are averaged to obtain an average transformation, which is subsequently inverted. **(5)** The inverted average transformation from step 4 is applied to the sharpened average intensity image from step 3. This step updates the template's shape to more closely reflect the average shape of the individual brains. The updated template becomes the new registration target for the next iteration of the SyGN algorithm. Note that the template construction process involves multiple iterations of this algorithm, progressing through several transform stages (Supplementary Figure 2).

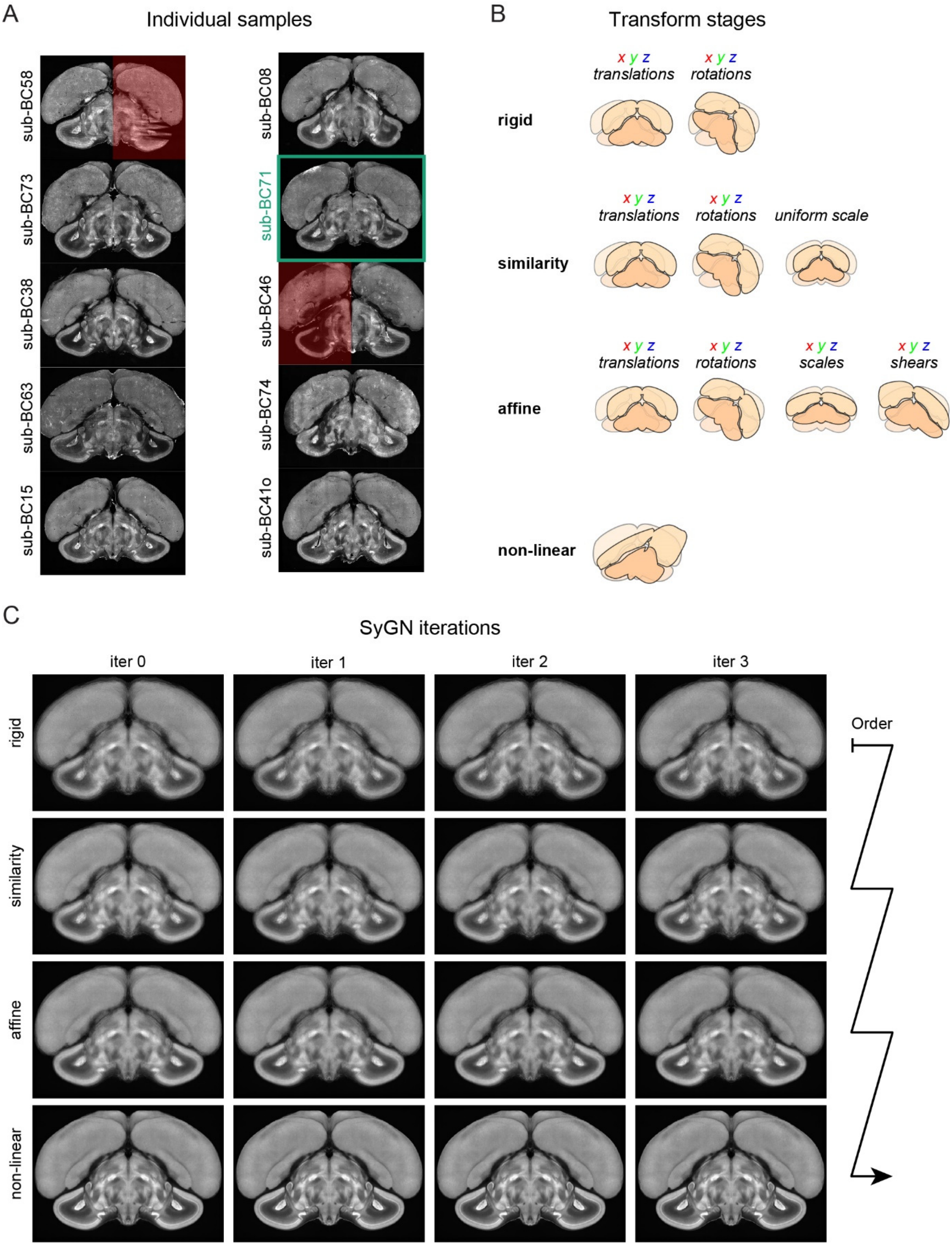


**Supplementary Figure 2: Average template construction through progressively higher-order transformations. (A)** Coronal slice views of 10 blackcap brains sectioned using STP tomography. Identifiers of individual animals displayed vertically. Red highlighted hemispheres were excluded from further processing steps due to tissue damage. The green highlighted brain (sub-BC71) is the example brain displayed in Figure 3. **(B)** Schematic depiction of the transformation types allowed at each of the four transform stages: rigid (3 translations + 3 rotations), similarity (rigid + 1 uniform scale), affine (rigid + 3 scales + 3 shears), non-linear symmetric diffeomorphic. **(C)** Coronal slice views of the average templates produced at each transform stage (rows) and SyGN iteration (columns), with four SyGN iterations per transform stage. The transformation parameters produced at each stage are used to initialize subsequent stages. The average template becomes progressively sharper (arrow indicates the order of execution), with the bottom right image corresponding to the final average template—i.e., the reference image of the atlas.
